# An assessment of efficacy of Iodine complex (Renessans) against SARS-CoV-2 in non-human primates (*Rhesus macaque*)

**DOI:** 10.1101/2020.11.17.377432

**Authors:** Muhammad Nawaz, Muhammad Asad Ali, Muhammad Adnan Ashraf, Muhammad Zubair Shabbir, Muhammad Abu Bakr Shabbir, Imran Altaf, Sohail Raza, Saira Rafique, Sohail Hassan, Nageen Sardar, Adnan Mehmood, Muhammad Waqar Aziz, Sehar Fazal, Muhammad Tahir Khan, Muhammad Moavia Attique, Ali Asif, Zia Ullah, Mubashir Iqbal, Talha Imtiaz, Muhammad Anwar, Nadia Mukhtar, Tahir Yaqub

**Affiliations:** Institute of Microbiology, University of Veterinary and Animal Sciences, Lahore, Pakistan; Institute of Public Health, Lahore, Pakistan

**Author notes:** Corresponding author Prof. Dr. Tahir Yaqub, Email ID.

**Keywords:** Iodine complex, Renessans, COVID-19, SARS-CoV-2, Rhesus macaque

## Abstract

Renessans is an iodine complex which has proven *in vitro* antiviral activity including Anti-SARS-CoV-2 activity. The present study was designed to determine its efficacy against SARS-CoV-2 in monkeys (*Rhesus macaque*). A total of 14 monkeys were divided into four groups: A) Prophylactic group (n=03), (B) Treatment group (n=03), (C) infection control group (n=04) and (D) negative control group (n=04) and were housed in BSL-3 Animal facility while group D was housed at another animal house. Group A was administered with Renessans @ 2.85 mg/7 kg from 5 days prior to the infection to 08 days post infections (DPI). Group B was administered with Renessans from 03-08 DPI @ 2.85 mg/7 kg. Group C was administered with WIF only. The infection @ 2 × 10^6^ TCID of SARS-CoV-2 was given to all group monkeys through intranasal and oral route under anesthesia. Nasal swab samples (at different times) and fecal matter on daily basis were collected for the detection of SARS-CoV-2 through real-time quantitative PCR. Three monkeys (one from each of group A, B and C) were euthanized at 07 DPI to determine the gross pathological lesions and SARS-CoV-2 detection from internal tissues. Nasal swabs from all the monkeys from group A, B and C were positive for SARS-CoV-2 at 02 and 07 DPI (Day 05 of treatment). At 14 DPI, all (100%) nasal swabs from group A were negative for SARS-CoV-2 while 50% and 100% were positive from group B and C, respectively. At 21 DPI, monkeys from group B were negative and all in group C were still positive for SARS-CoV-2. Similarly, fecal matter of monkeys in group A and B was returned negative in significantly lesser time as compared to monkeys from infection control group. Based on these research findings it is concluded that the Renessans has *in-vivo* SARS-CoV-2 activity and may result in early clearance of SARS-CoV-2. Therefore, a clinical trial of the drug in COVID-19 patients may reveal its anti-COVID-19 potential.

## Introduction

Severe acute respiratory syndrome-related coronavirus-2 (SARS-CoV-2) was first time reported as the etiologic agent of coronavirus disease 2019 (COVID-19) in December 2019 at a wholesale seafood market in Wuhan, Hubei province, China (Lake, 2020; WHO, 2020). According to the World Health Organization (WHO) that more than 5.3 million confirmed cases and around 340,000 fatalities have been reported all over the world since its first report (WHO, 2020).

SARS-CoV-2 belongs to the Coronaviridae family and *Nidovirales* order. Coronaviruses are divided into alpha (α), beta (β), gamma (γ) and delta (δ) groups. This virus is from β-coronaviruses (Muniyappa and Gubbi, 2020; Yi et al, 2020; Ye et al., 2020). SARS-CoV-2 is a single stranded non-segmented positive sense RNA with a size of 30 kb. The genome contains sequences for replicases, papain like proteases, endoribonuclease, and spike proteins. It is important to note that spike proteins of SARS-CoV-2 are not alike from those of SARS-CoV (Tang et al., 2020; Hoffmann et al., 2020). SARS-CoV-2 is round in shape and has an envelope. Spike proteins (S1 and S2) and glycoproteins are present on its envelope. These spike proteins bind with Angiotensin-Converting Enzyme-2 (ACE-2) receptors of host cells and helps the virus to enter the cell by endocytosis. While, the membrane protein (M) of envelop determines the virus shape (Tang et al., 2020).

SARS-CoV-2 can spread from one human to another human through coughing and sneezing. Predilection site of the virus is lung’s alveolar epithelial type 2 (AT2) cells. Several studies reported that the spike proteins of SARS-CoV-2 bind to ACE-2 receptors present on AT2 cells (Wang et al., 2020; Li et al., 2019). It has been reported that ACE-2 receptors also present on tubular epithelium of kidney, pancreas, heart, and endothelial cells (Diao et al., 2020; Liu et al., 2020; Zheng et al., 2020). Upon entering into the host cell, the virus releases its positive sense RNA that dictates host cell machinery and produce new virions (Sigrist et al., 2020).

SARS-CoV-2 infection can be asymptomatic and in most cases may cause mild to severe complications (Cao, 2020). Given the prevalence of asymptomatic individuals and limited availability of molecular testing in different parts of world, it is believed that true number of infections may be several fold higher than the estimates of WHO (Cheng et al., 2020).

Noteworthy, therapeutic options for SARS-CoV-2 have not been developed so far and hence only supportive therapy is provided to the patients (Raza et al., 2020). Therefore, present study was designed to develop a treatment for SARS-CoV-2. This study was based on our previous *in vitro* study (under review) findings in which Renessans (antiviral drug) showed promising results. In current study, we determine its *in vivo* efficacy against SARS-CoV-2 in monkeys (*Rhesus macaque*). A total of 14 monkeys were divided into 4 groups and SARS-CoV-2 infection was given to group A, B and C. Pre and post infection nasal as well as fecal sampling was performed for the detection of SARS-CoV-2 by real-time quantitative PCR. Furthermore, one monkey from each group A, B and C were euthanized for determining the gross pathological lesions as well as SARS-CoV-2 from different tissues samples. Present study findings did reveal that Renessans have antiviral activity and helps in early clearance of SARS-CoV-2. We believe that current study findings will provide a baseline for clinical trial against SARS-CoV-2 infection and hence helps in the development of therapeutic option for SARS-CoV-2 infection.

## Materials and Methods

### Experimental Design

A total of 14 monkeys (*Rhesus macaque*) were obtained from wildlife department of Pakistan to determine the *in vivo* efficacy of antiviral drug (Renessans) against SARS-CoV-2. Monkeys were weighted and divided into four groups: A) Prophylactic group (n=03), (B) Treatment group (n=03), (C) infection control group (n=04) and (D) negative control group (n=04). These non-human primates were housed in Animal Biosafety Laboratory-3 (ABSL-3) of Institute of Microbiology, University of Veterinary and Animal Sciences (UVAS) Lahore, Pakistan for one month under standard conditions of ambient temperature (22 ± 2 °C). Food and water was provided to monkeys *ad libitum* throughout the experiment.

### Infection

Before starting the experiment, approval was taken from Institutional Biosafety committee (IBC) of UVAS, Lahore, Pakistan. Furthermore, experiment was performed according to the ethical guideline of UVAS, Lahore, Pakistan.

The antiviral drug (Renessans) was administered @ 2.85 mg/7 kg at the date 22 August 2020 to group A from 5 days prior to the infection to 08 days post infection (DPI). Group B was administered with Renessans after the onset of clinical signs and symptoms from 03-08 DPI @ 2.85 mg/7 kg. Group C was administered with WIF only. SARS-CoV-2 (GenBank accession number MW031802) infection @ 2 × 10^6^ TCID was given to group A, B and C through intranasal and oral route under anesthesia (mixture of ketamine and xylaz) at the date 26 August 2020. Additionally, body temperature of group A, B and C monkeys was also monitored on daily basis throughout the experiment after the onset of clinical signs and symptoms.

### Fecal and Nasal swab Sampling

Fecal and nasal swab sampling was performed to determine the shedding of SARS-CoV-2 through these routes. All monkeys of group A, B and C were anesthetized for nasal sampling. We did nasal sampling five times during the whole experiment; firstly one day before the infection and then at 2 DPI, 7 DPI, 14 DPI and 21 DPI from all monkeys of group A, B and C. However, we started fecal sampling on daily basis from day 0 (infection date 26-08-2020) to 16-09-2020 (experiment ending date). For better understanding, experimental plan or design is given in Figure 1.

**Figure 1:**
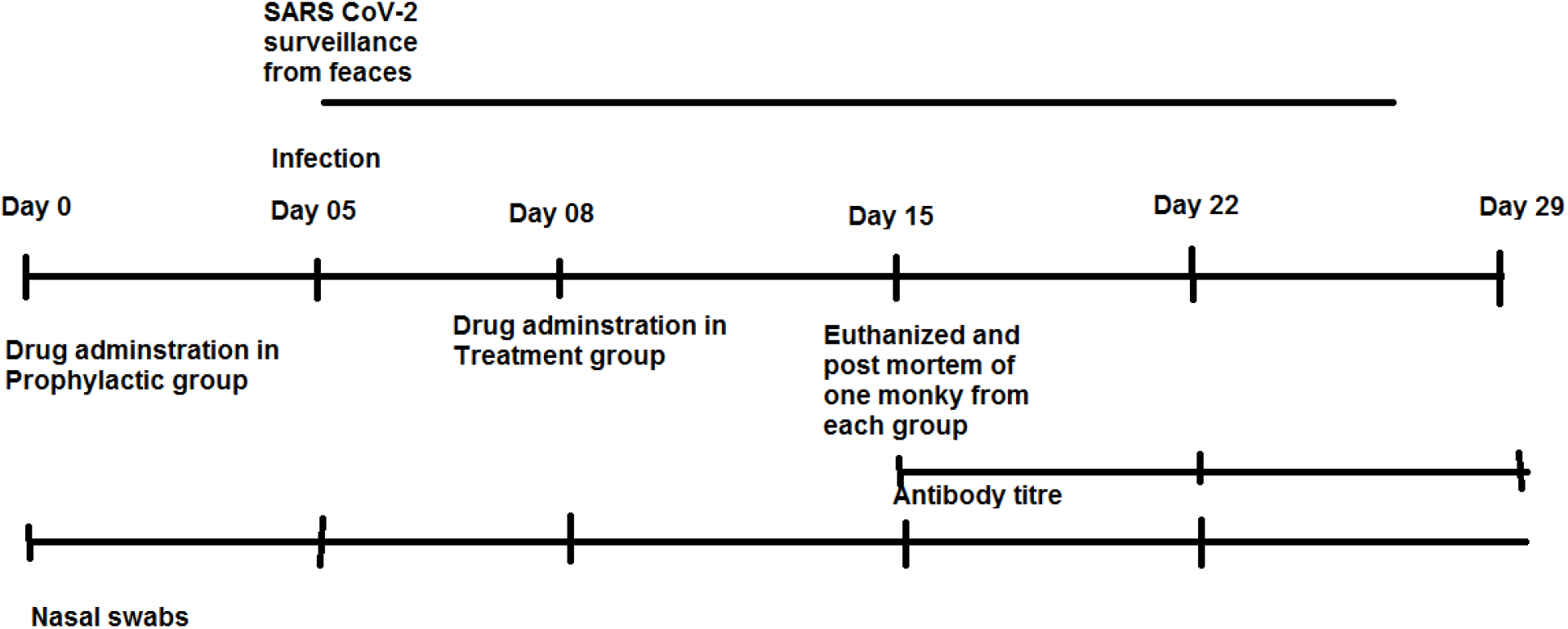
Detailed experimental plan for determining the *in vivo* efficacy of antiviral drug (Renessans) against SARS-CoV-2.

### SARS-CoV-2 detection from tissues by real time quantitative PCR

A total of 3 monkeys (one from each group A, B and C) were euthanized for the determination of gross pathological lesions and SARS-CoV-2 from different tissues by real-time quantitative PCR. These monkeys were euthanized by giving the intra-cardiac injection of potassium chloride (Kcl) @ 10 mL. Gross pathological lesions were noted upon postmortem and tissue samples were taken and stored at −80 °C till further use.

## RESULTS

### 1. SARS-CoV-2 detection from fecal samples of Monkeys by Real-time Quantitative PCR

Fecal samples were collected from A, B and C group monkeys at different times for the detection of SARS-CoV-2 by real time qualitative RT PCR. At 7 DPI and 14 DPI, all group C monkeys were found positive for SARS-CoV-2 while the said virus was detected from fecal matter of A (P2) and B (T3) group monkeys (one each). Noteworthy; at 21 DPI, our A group monkey P2 found negative for SARS-CoV-2 and group B (T3) monkey still found positive for SARS-CoV-2. All monkeys of group C were still shedding the virus. We can also say that fecal matter of monkeys in prophylactic group (P2 monkey) and treatment (T3 monkey) was returned negative in significantly lesser time as compared to monkeys from infection control group (Table 1). These findings suggesting that antiviral drug (Renessans) did have *in-vivo* SARS-CoV-2 activity and may result in early clearance of SARS-CoV-2.

**Table 1:**
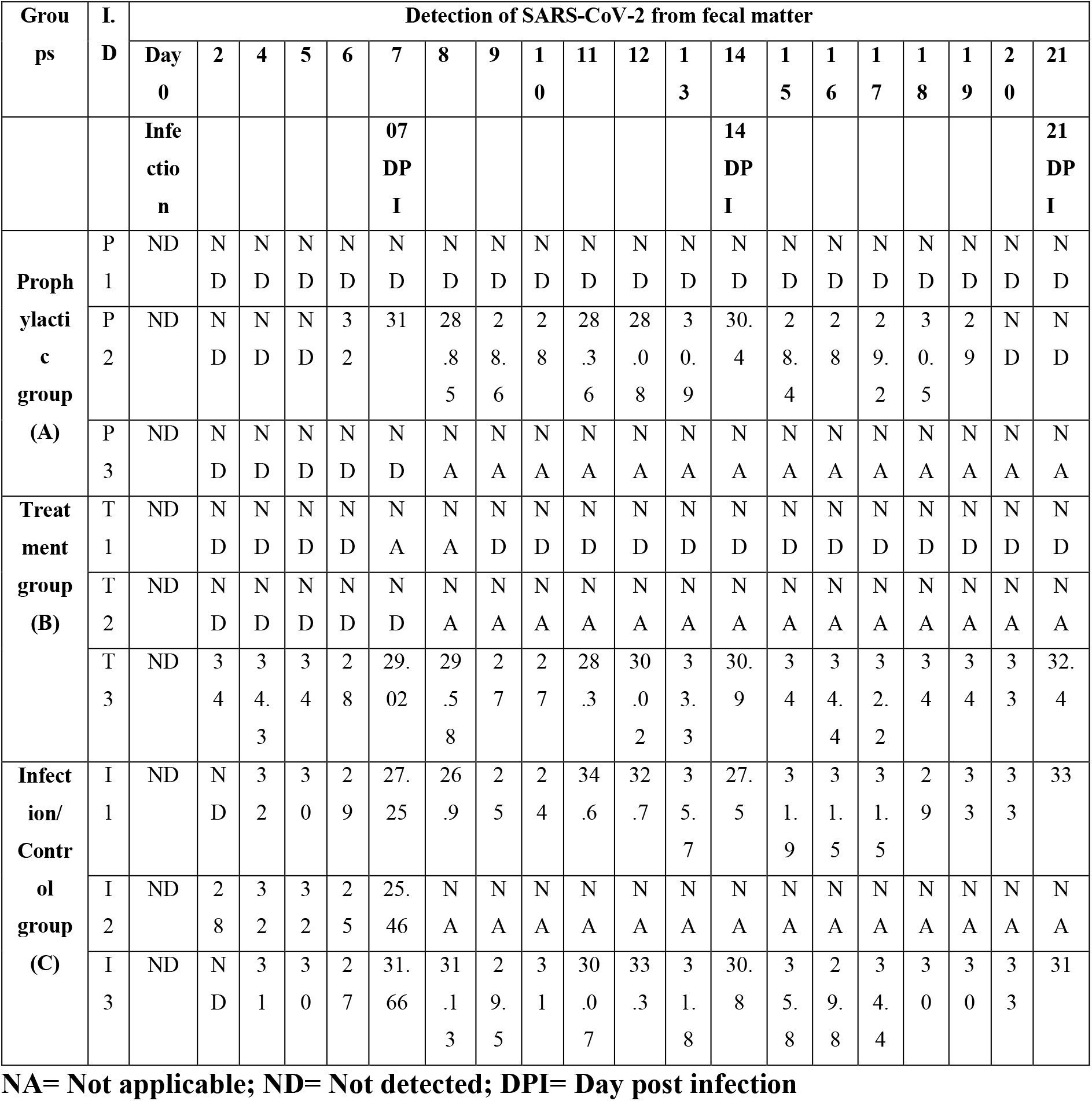
Detection of SARS-CoV-2 from fecal samples of Monkeys by Real-time Quantitative PCR.

### 2. SARS-CoV-2 detection from nasal swab samples of Monkeys by Real-time Quantitative PCR

Nasal swabs were collected from A, B and C group at different times for the detection of SARS-CoV-2 by real time qualitative RT PCR. Pre-infection nasal swab sampling was also formed to detect the SARS-CoV-2 by real-time quantitative PCR from all the monkeys of group A, B and C and we found that all monkeys of these three groups were negative for SARS-CoV-2. However; after the 48 hours of infection, all the monkeys from group A, B and C were found positive for SARS-CoV-2 at 02 and 07 DPI. Interestingly, all (100%) nasal swabs from group A and B were negative for SARS-CoV-2 at 14 and 21 DPI. However, monkeys of C group were still found positive for SARS-CoV-2 at 14 and 21 DPI. Based on these findings we can say that Renessans (antiviral drug) did have positive effect and helps in the early recovery of group A and B monkeys from SARS-CoV-2. Detailed results are given in Table 2.

**Table 2:** Detection of SARS-CoV-2 from nasal swab samples of Monkeys by Real-time Quantitative PCR.

| Groups | Monkey I.D | Detection of SARS-CoV-2 from nasal swab |  |  |  |  |  |
| --- | --- | --- | --- | --- | --- | --- | --- |
|  |  | Pre-infection | 26/8<br>0 Day | 2DP1 | 7DPI | 14 DPI | 21 DPI |
| Prophylactic (A group) | P1 | ND | Infection<br>date | 21 | 25.16 | ND | ND |
|  | P2 | ND |  | 28 | 36 | ND | ND |
|  | P3 | ND |  | 18 | 28.37 | NA | NA |
| Treatment (B group) | T1 | ND |  | 18 | 25 | ND | ND |
|  | T2 | ND |  | 24 | 27.42 | NA | NA |
|  | T3 | ND |  | 26 | 25 | ND | ND |
| Infection (C group) | I1 | ND |  | 21 | 25.2 | 33 | 32 |
|  | I2 | ND |  | 19 | 28.7 | NA | NA |
|  | I3 | ND |  | 17 | 27.27 | 26.8 | 29 |
|  | I4 | ND |  | 24 | NA | NA | NA |

NA= Not applicable; ND= Not detected; DPI= Day post infection

### 3. SARS-CoV-2 detection from different tissues of group A, B and C monkeys by Real-time Quantitative PCR

To determine the gross pathological lesions during the SARS-CoV-2 infection, one monkey (3C) from group C was euthanized at 02 DPI while three monkeys (one from each of group A, B and C) were euthanized at 07 DPI. Gross pathological lesions were noted and SARS-CoV-2 detection from different tissues was performed by real-time quantitative PCR. Upon real-time PCR, SARS-CoV-2 was detected from internal tissue i.e intestine, lung, heart and spleen. At 07 DPI, lung, trachea and heart tissues of monkeys in the infection control group (group C) were positive for SARS-CoV-2 while lung, trachea, heart tissues of monkeys from group A and B were negative. These findings also suggested that the antiviral drug Renessans did have positive effect in SARS-CoV-2 infection. Detailed results are given in table 3.

**Table 3:**
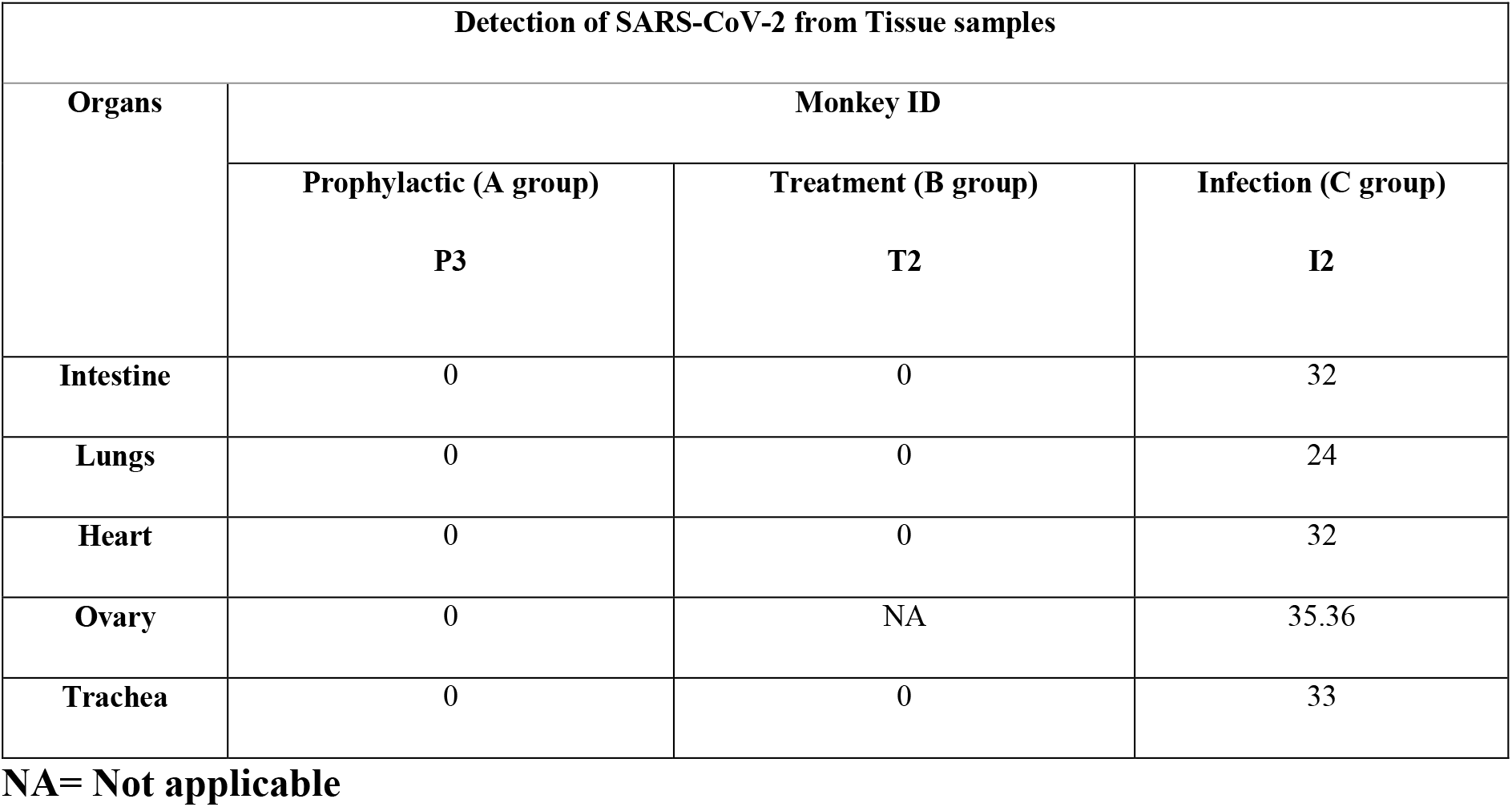
Detection of SARS-CoV-2 from different tissues of group A, B and C monkeys by Real-time Quantitative PCR.

| Detection of SARS-CoV-2 from Tissue samples |  |  |  |
| --- | --- | --- | --- |
| Organs | Monkey ID |  |  |
|  | Prophylactic (A group) | Treatment (B group) | Infection (C group) |
|  | P3 | T2 | I2 |
| Intestine | 0 | 0 | 32 |
| Lungs | 0 | 0 | 24 |
| Heart | 0 | 0 | 32 |
| Ovary | 0 | NA | 35.36 |
| Trachea | 0 | 0 | 33 |

NA= Not applicable

## Discussion

Present study was designed to determine the *in-vivo* efficacy of Renessans (antiviral drug) in non-human primates (*Rhesus macaque)* against SARS-CoV-2. We observed that Renessans antiviral drug showed promising results against SARS-CoV-2 in *Rhesus macaque*. We start collecting the fecal samples at 0 day of infection (SARS-CoV-2) on daily basis till the end of *in-vivo* experiment. Real-time quantitative PCR was used to detect SARS-CoV-2 and it was observed that at 7 DPI and 14 DPI, only one monkey in each group A (P2) and B (T3) were positive for SARS-CoV-2 while all monkeys of group C positive for said virus. At 21 DPI, group A monkey P2 found negative for SARS-CoV-2 while group B monkey T3 still found positive for SARS-CoV-2. However, monkeys from group C were still shedding the virus. Similarly, pre-infection nasal swab sampling was performed from all group monkey i.e. A, B and C for SARS-CoV-2 infection by real time quantitative PCR and it was observed that monkeys of these groups were negative. However; at 02 DPI and 07 DPI, all the monkeys from group A, B and C were found positive for SARS-CoV-2. Noteworthy; at 14 DPI and 21 DPI, nasal swab samples of prophylactic (A) and treatment (B) group were found negative for SARS-CoV-2. However, monkeys of C group were still found positive for SARS-CoV-2 at 14 and 21 DPI. This might be due to positive effect of Renessans against SARS-CoV-2 and it may also possible that prophylactic (A) and treatment (B) group were recovered early because of SARS-CoV-2 specific immune response. Present study findings are in line with recently published study findings (Kuri et al., 2020; Mathew et al., 2020).

The monkeys of group A, B and C were infected @ 2 × 10^6^ TCID of SARS-CoV-2 through intranasal and oral route under anesthesia. Nasopharyngeal swab sampling is a standard method to detect SARS-CoV-2 (WHO, 2020), and hence pre and post infection nasal swab sampling of group A, B and C was performed and SARS-CoV-2 was detected by real time quantitative PCR. At 2 and 7 DPI, all the monkeys from group A, B and C were found positive for SARS-CoV-2; however, it is interesting to note that all (100%) nasal swabs from group A and B were negative for SARS-CoV-2 at 14 and 21 DPI. Understandably, monkeys of C group were still found positive for SARS-CoV-2 at 14 and 21 DPI. Based on these findings we can say that Renessans (antiviral drug) did have positive effect and helps in the early recovery of group A and B monkeys from SARS-CoV-2.

Gross pathological lesions as well as presence of SARS-CoV-2 virus was determined by euthanizing the one monkey in each group A, B and C at 07 DPI. Upon real time quantitative PCR of postmortem biopsy samples, SARS-CoV-2 virus was detected from lungs, spleen, intestine and heart. This suggest that SARS-CoV-2 can infect other organs apart from lungs. Similar findings were also observed in another study where they detected SARS-CoV-2 by RT-PCR in heart and liver (Tian et al., 2019). However, it is important to note that gross pathological lesion were less severe in group A and B than the group C, suggesting that Renessans did have antiviral activity and helps in the early recovery of SARS-CoV-2 infected monkeys.

## Conclusion

In the light of current study findings, it is concluded that the Renessans has an *in-vivo* SARS-CoV-2 activity and may result in early clearance of SARS-CoV-2. Therefore, we believe that current study may provide a basis for clinical trial of the drug in SARS-CoV-2 patients and reveal its anti-SARS-CoV-2 potential.

